# Statistical evaluation of metaproteomics and 16s rRNA amplicon sequencing techniques for the study of the gut microbiota establishment of infants with cystic fibrosis

**DOI:** 10.1101/2022.04.18.488713

**Authors:** Claudia Saralegui, Carmen García-Durán, Eduardo Romeu, María Luisa Hernáez-Sánchez, Ainhize Maruri, Natalia Bastón-Paz, Adelaida Lamas, Saioa Vicente, Estela Pérez-Ruiz, Isabel Delgado, Carmen Luna-Paredes, Juan de Dios Caballero, Javier Zamora, Lucía Monteoliva, Concepción Gil, Rosa del Campo

**Author notes:** **Correspondence:** Departamento de Microbiología y Parasitología, Universidad Complutense de Madrid, Spain, and Servicio de Microbiología, Hospital Universitario Ramón y Cajal, Madrid, Spain. Both authors contributed equally (Claudia Saralegui, Carmen García-Durán).

## Abstract

The newborn screening for cystic fibrosis (CF) can identify affected but still asymptomatic infants. The selection of omic technique for gut microbiota study is therefore crucial due to both the small amount of feces available and the low microorganism load. Our aim was to compare the agreement between 16S rRNA amplicon sequencing and metaproteomics by a robust statistical analysis including McNemar (taxa presence) test and Bland–Altman (taxa abundance) data plotting for studying the sequential establishment of the gut microbiota during the first year of life in 8 infants with CF (28 fecal samples). The taxonomic assignation was similar by the two techniques, whereas certain discrepancies were observed in the abundance detection, mostly the underrepresentation of *Bifidobacterium* and the overestimation of certain *Firmicutes* and *Proteobacteria* by amplicon sequencing. During the first months of life, the CF gut microbiota is characterized by poor alpha diversity, a significant enrichment of *Ruminococcus gnavus*, the expression of certain virulent bacterial traits, and the detection of human inflammation-related proteins. Our study provides an extended comparative analysis with robust statistical support that could optimize the use of both approaches for gut microbiota research. Metaproteomics provides information on composition and functionality, as well as data on host-microbiome interactions. Its strength is the identification and quantification of *Actinobacteria* and certain classes of *Firmicutes*. Both techniques detected an aberrant microbiota in infants with CF during their first year of life, dominated by the enrichment of *R. gnavus* within a human inflammatory environment.

## INTRODUCTION

The establishment of the microbiota at birth affects human health during the rest of life (1). In cystic fibrosis (CF), a genetic disease diagnosed by neonatal screening, the colonization of the lung airways by pathogenic bacteria impacts survival (2). The life expectancy of individuals with CF has increased considerably in recent years, and comorbidities beyond those of the respiratory tract have been revealed, mostly related to the digestive tract (3). The metabolic activity of the gut microbiota has a direct impact on nutritional status; in subjects with CF, this activity has a major effect on their infectious lung disease (4).

The gut microbiota in CF is conditioned by the following factors: transmembrane regulator dysfunction, which creates a thick mucus layer; the acidic environment caused by impaired bicarbonate secretion; maldigestion; malabsorption; steatorrhea due to pancreatic insufficiency; and the high intake of antibiotics for pulmonary exacerbations (5, 6). Significant differences in the microbiota’s bacterial composition and alpha diversity have been reported compared with healthy infants (7-9).

Currently, 16S rRNA amplicon sequencing is the most extensive, rapid, and cost-effective technique for characterizing complex bacterial communities, such as the gut microbiota (10). The technique’s main weakness results from the variable affinity of the primers for flanking regions. Overrepresentation has been found for certain genera, whereas others, mainly *Bifidobacterium*, are consistently underrepresented (11). Even with optimized primers, amplicon sequencing often provides deficient taxonomic assignment at the genus and species levels due to the short length of the amplicons. On the other hand, metaproteomics applies high-resolution mass spectrometry (MS) in the detection and quantification of proteins, allowing both taxonomic identification and functional characterization of all present microorganisms, not just bacteria. The recent introduction of updated bioinformatics tools, such as MetaLab, has considerably improved the data analysis (12); for lesser-represented species, however, the analysis remains inaccurate (13). Available data suggest that only 10%-20% of the total protein content of a clinical sample is identified, corresponding to only one-third of the entire microbiota. Additional strategies have recently been suggested as prefractionation steps to circumvent instrument sensitivity problems and access low-abundance proteins of biological importance (14). The metaproteomic strengths include a high resolution in bacterial species identification, the overview of functional and metabolic activity of microorganisms, and most notably, the detection of host proteins, expanding the knowledge of host-microbiome interactions (15, 16), which is particularly interesting in CF (17).

The combination of amplicon sequencing and metaproteomics techniques provides a wider characterization of microbial communities and their interaction with the host (18-20), and the integration of the results could support accurate predictions (21). However, few studies have included a statistical analysis comparing the results by both methodologies, especially of challenging samples such as those from neonates, characterized by limited sample quantity and potentially low bacterial load. The aim of this study was to perform a robust statistical analysis on the concordance of amplicon sequencing and metaproteomics for the study of the sequential establishment of the gut microbiota of a small representative cohort of infants with CF.

## RESULTS

### Statistical concordance of the results of amplicon sequencing and metaproteomics

We analyzed the fecal samples (n=28) using both techniques for the bacterial identification (presence/absence) and quantification (abundance), then respectively applied the McNemar and Bland–Altman/Wilcoxon tests to detect significant discrepancies (Figure 1 and Additional Files 3 and 4). Globally, 685 ASVs (9 phyla, 13 classes, 130 genera, and 90 species) were identified by amplicon sequencing, and 317 taxa (8 phyla, 17 classes, 65 genera, and 133 species) were identified by metaproteomics.

**FIGURE 1.**
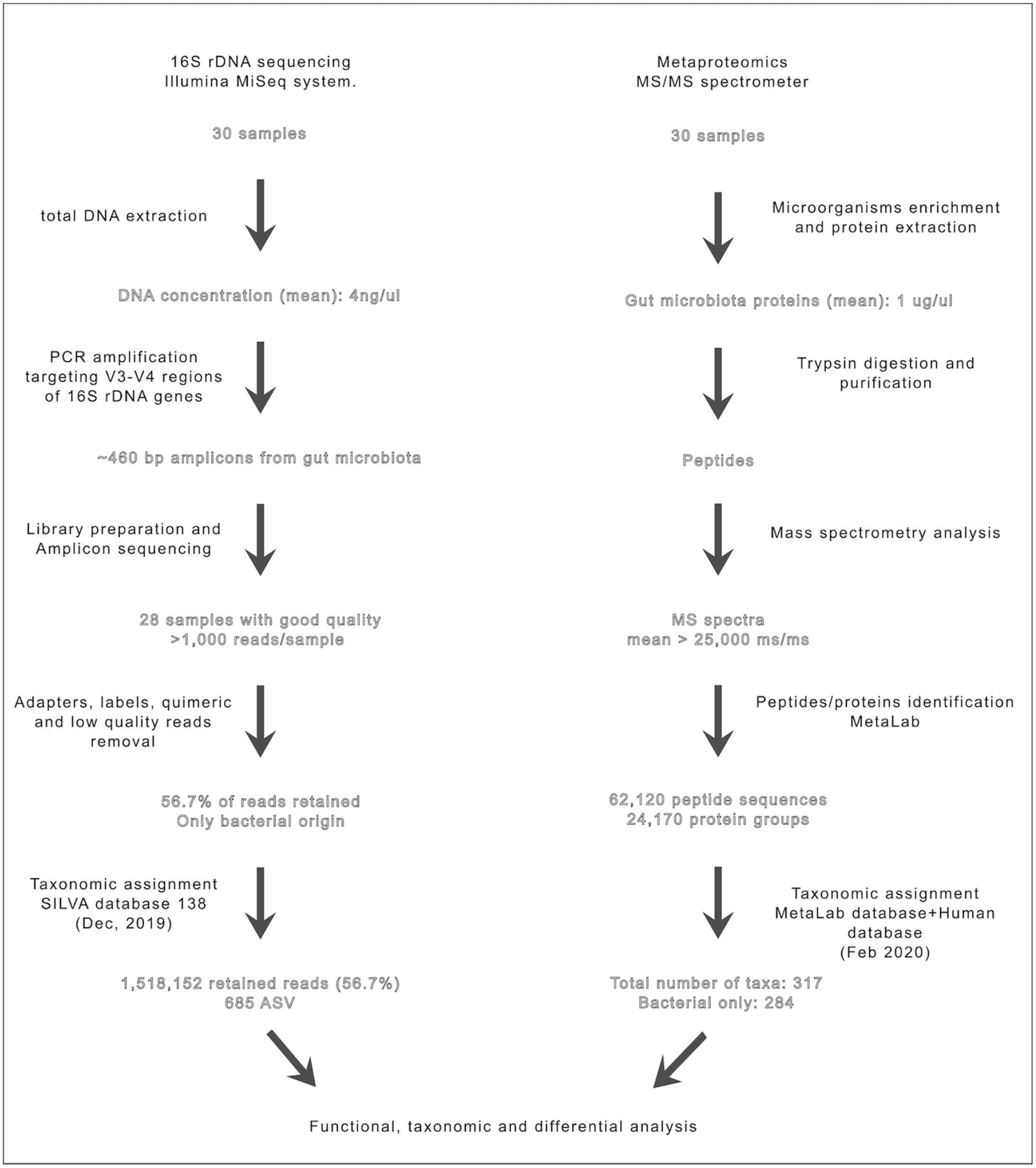
Amplicon sequencing and metaproteomics overall workflow for microbiota compositional and functional analysis together with the main experimental results output from each step.

Minor statistical discrepancies were observed in taxa presence/absence, most of which were related to the species level (Figure 2, Table 2, and Additional File 5). Furthermore, we detected certain nominal inconsistencies related to the taxonomic classification by each database, such as the inclusion of the *Tenericutes* phylum from metaproteomics in the *Firmicutes* phylum according to amplicon sequencing and the recent taxonomic reclassification of *Proteobacteria* classes (Table 2).

**Table 1.**
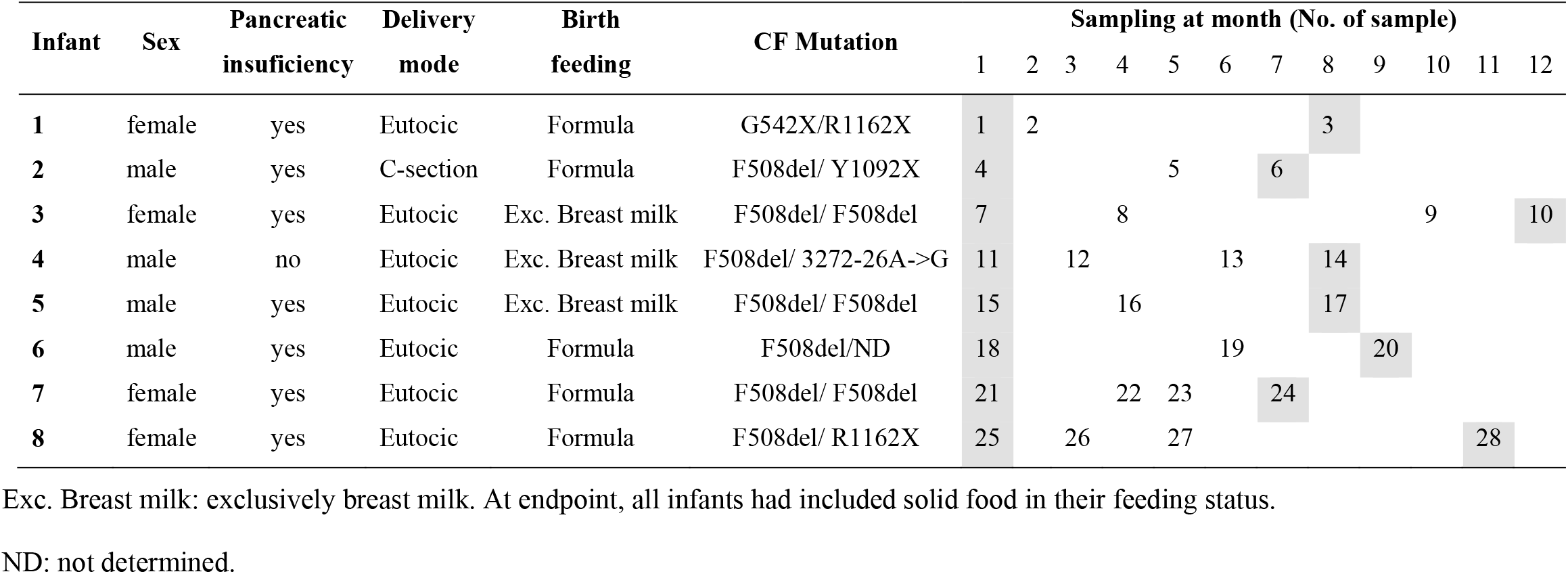
Relevant clinical characteristics of infants and sampling. All samples (n=28) were included in the methodological analysis, whereas samples in grey (n=16, the initial 8 and evolved 8 last samples per infant) were considered to analyze gut microbiota establishment analysis.

**Table 2.**
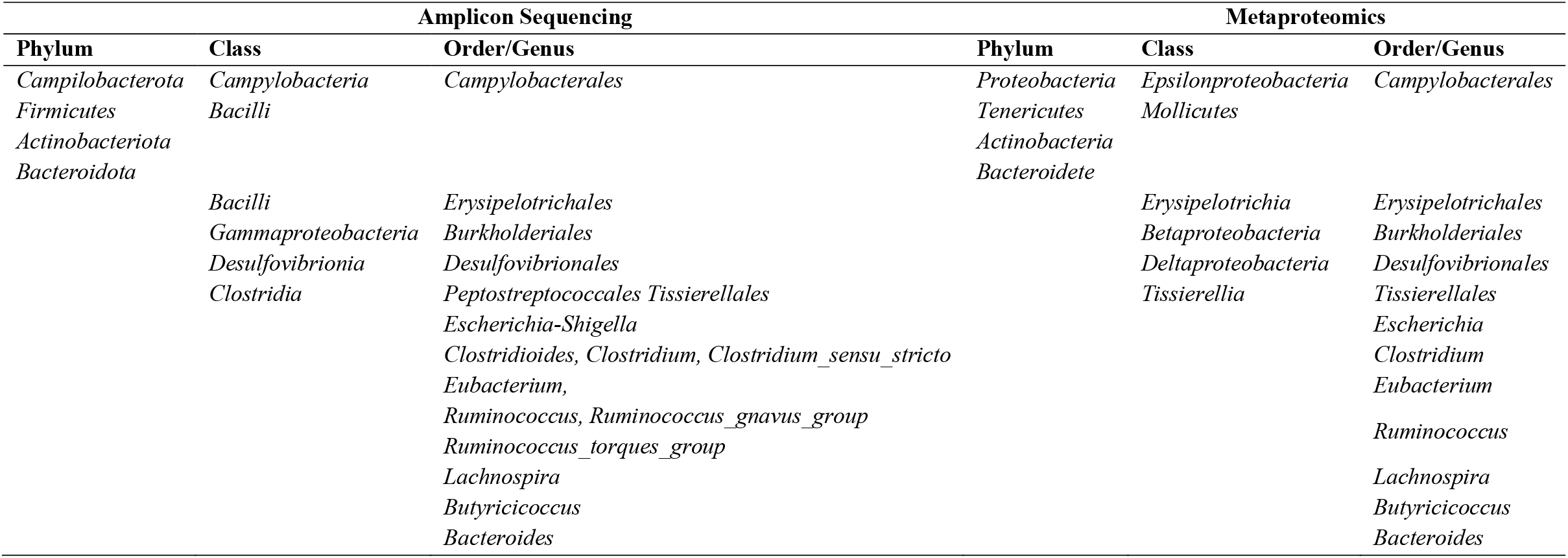
Incongruencies in taxonomic labeling or bacteria misclassification at different ranks by amplicon sequencing and metaproteomics.

**FIGURE 2.**
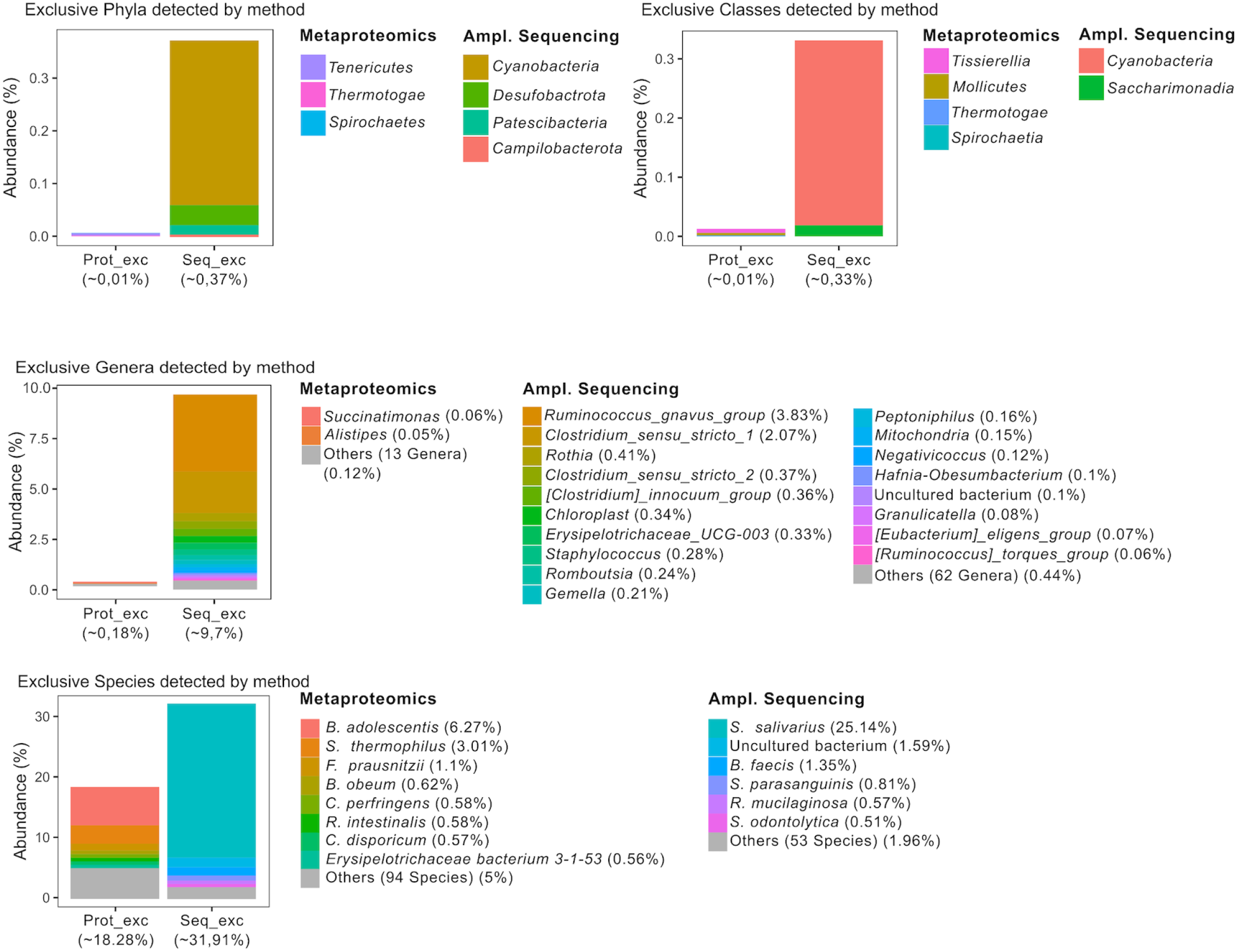
Exclusive taxa and their relative abundance detected by amplicon sequencing and metaproteomics. Twenty-eight samples were included in the analysis and each relative abundance was normalized per technology. Amplicon sequencing does not have enough resolution for taxonomic annotation at species rank even though this information was given.

Figure 3 summarizes the results of the taxa abundance analysis by the Bland– Altman test combined with the Wilcoxon test. Overrepresentation of the *Actinobacteriota* (*Actinobacteria* class and, specifically, *Bifidobacterium* genus) and *Clostridia* class (*Clostridioides* genus) were detected when metaproteomics was employed, whereas in amplicon sequencing, the overrepresented taxa were *Firmicutes* (*Bacilli* class, *Enterococcus, Streptococcus*, and *Ruminococcus* genera) and *Proteobacteria* (*Gammaproteobacteria* class and, specifically, *Escherichia-Shigella* genus).

**FIGURE 3.**
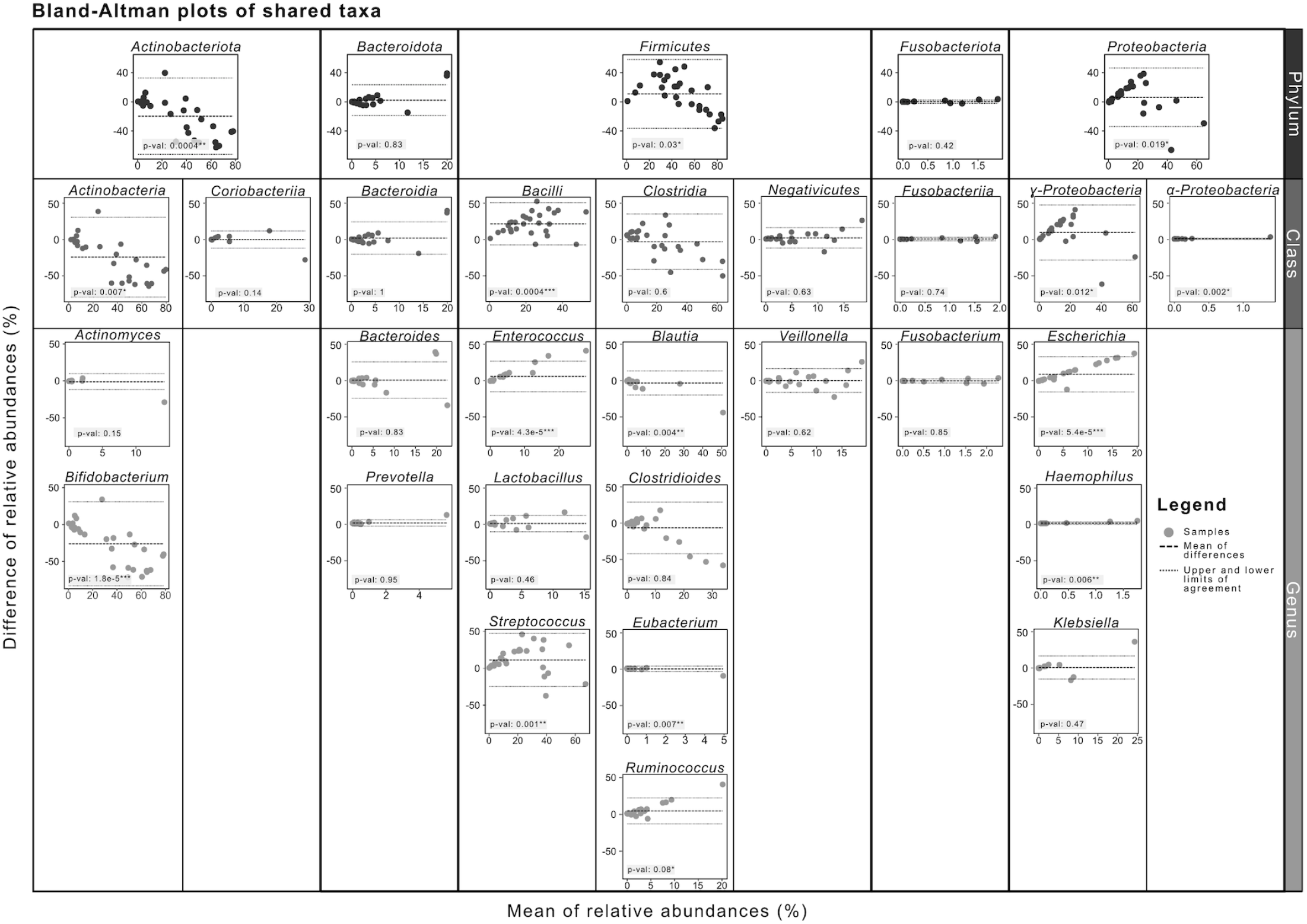
Bland-Altman plots representing mean of relative abundance (%) of each taxa per sample and technology against difference of relative abundance (amplicon sequencing data minus metaproteomics data) of each taxa per sample and technology (%). P-value from nonparametric Wilcoxon rank-sum paired test is included in the plots. ***: p<0.001; **: p<0.01; *: p<0.05.

### Bacterial establishment in the CF gut

We assessed the dynamics of bacterial composition during the first year of life in the initial (n=8) and evolved (n=8) samples of each infant by amplicon sequencing (192 ASVs) and metaproteomics (92 taxa). Unexpectedly, the alpha diversity indexes were higher in the initial samples, although statistical significance was only reached at the class level in amplicon sequencing (1.4±0.2 vs. 1.1±0.4, *p*=0.02) (Figure 4A).

**FIGURE 4.**
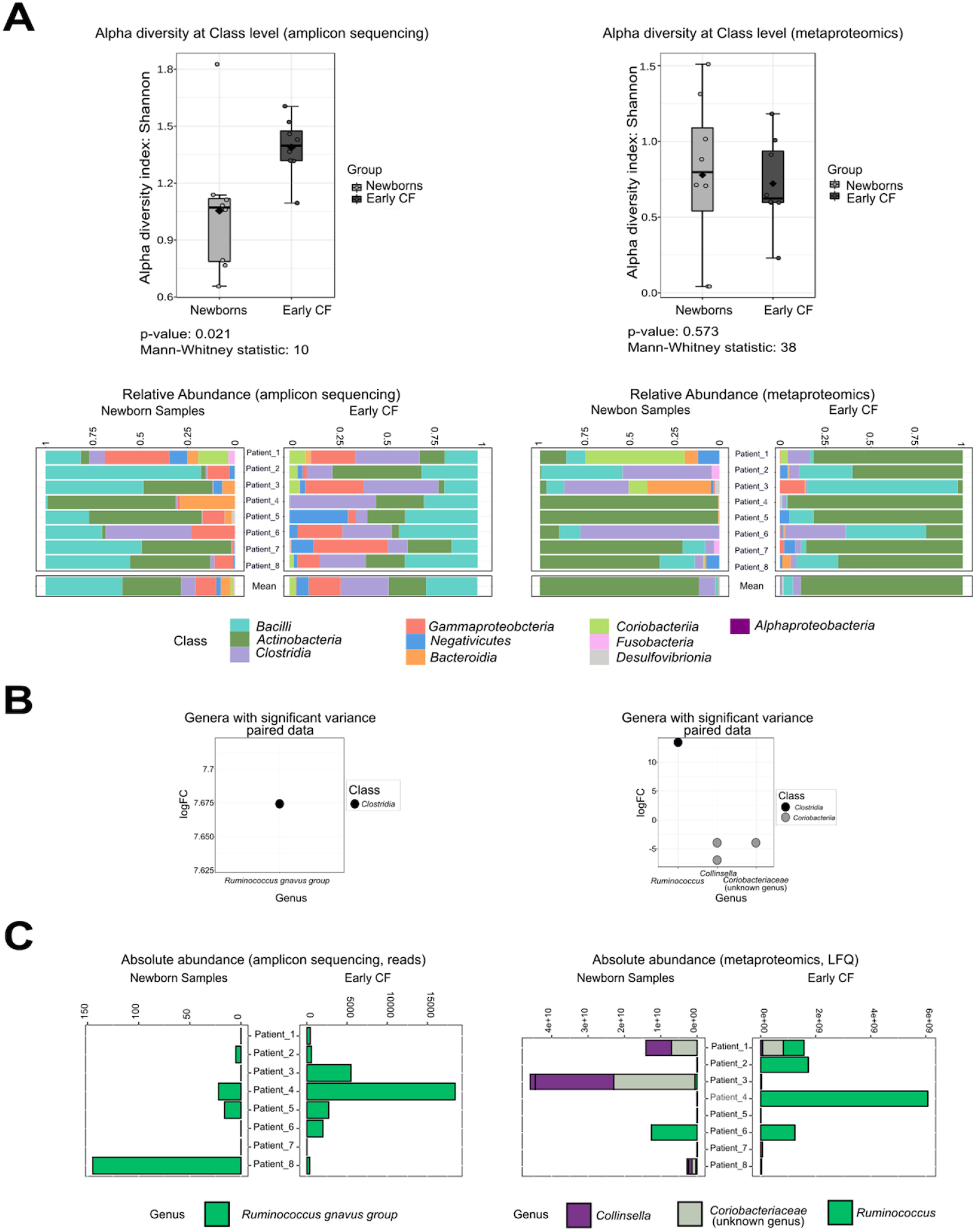
Significant differences in microbiota composition in CF infants during their first months of life inferred from amplicon sequencing (left panels) and metaproteomics (right panels). A: alpha diversity indexes in newborn and evolved CF samples (n=8) at Class level (top), and relative abundances of main bacterial classes detected in each infant at the beginning and at the end of the study (bottom). B: bacterial classes with significant fold-change shifts (increased in newborn samples: logFC>0; increased in early CF samples: logFC<0) between groups after adjusting for baseline differences among infants. C: absolute abundances of those significantly variant taxa in newborn and early CF groups. LFQ: intensity units from metaproteomics.

To decipher the relevant abundance fluctuations in the evolved samples, only the taxonomic traits with the greatest positive or negative variations were considered after discarding the initial abundance. Using amplicon sequencing, a significant abundance increase was detected only for the *Ruminococcus gnavus* group (log_2_ 7.7, *p*=0.0005) (Figure 4B). Metaproteomics failed to detect significant changes at the typical significance value (*p*<0.05). With less strict criteria (*p*<0.1), however, 4 OTUs assigned to the *Ruminococcus* and *Collinsella* genera had a significant fold-change shift (Figure 4B; right panel).

### Functional assignment by metaproteomics

Functional information on the detected proteins was obtained by MetaLab software, using the sum of the peptide intensities for each protein (total intensity of the protein). Both the number and intensity of the proteins in the initial (5718 and 4.86e12, respectively) and the evolved (7029 and 4.70e12, respectively) CF fecal samples were comparable. MetaLab assigned the functional profile by matching proteins with the Cluster of Orthologous Group (COG) database with a 97% successful assignment for name/category. A previous normalization process of the protein intensities (intensity of the function/sum of the intensities in that sample) was performed, and the 2-way analysis of variance detected 14 COGs (6 from the initial and 8 from the evolved samples) with statistical relevance (Table 4). However, there were no significant differences in the general functional profiles in the enrichment analysis, of which “energy production and conversion” (C), “carbohydrate transport and metabolism” (G), and “translation, ribosomal structure, and biogenesis” (J) were the COG categories with greater intensity (Figure 5A).

**Table 3.** Contingency table for McNemar test accounting for the number of samples where different bacteria (class and genus level) were or were identified by one, both or none of the technologies. Significance level<0.05.

| Genus | Number of samples detected by |  |  |  | *P* value |
| --- | --- | --- | --- | --- | --- |
|  | Amplicon sequencing only | Metaproteomics only | Both | None |  |
| Class |  |  |  |  |  |
| *Alphaproteobacteria* | 8 | 1 | 9 | 10 | 0.02 |
| Genus |  |  |  |  |  |
| *Eggerthella* | 12 | 0 | 1 | 15 | 0.001 |
| *Anaerococcus* | 11 | 0 | 1 | 16 | 0.002 |
| *Haemophilus* | 15 | 2 | 5 | 6 | 0.003 |
| *Coprococcus* | 1 | 10 | 1 | 16 | 0.007 |
| *Lachnoclostridium* | 0 | 9 | 2 | 17 | 0.007 |
| *Roseburia* | 0 | 9 | 2 | 17 | 0.007 |
| *Eubacterium* | 1 | 14 | 6 | 7 | 0.01 |
| *Enterobacter* | 10 | 1 | 3 | 14 | 0.01 |
| *Enterococcus* | 10 | 1 | 14 | 3 | 0.01 |
| *Escherichia* | 11 | 1 | 14 | 2 | 0.01 |
| *Actinomyces* | 14 | 4 | 6 | 4 | 0.03 |
| *Collinsella* | 8 | 1 | 5 | 14 | 0.04 |

**Table 4.**
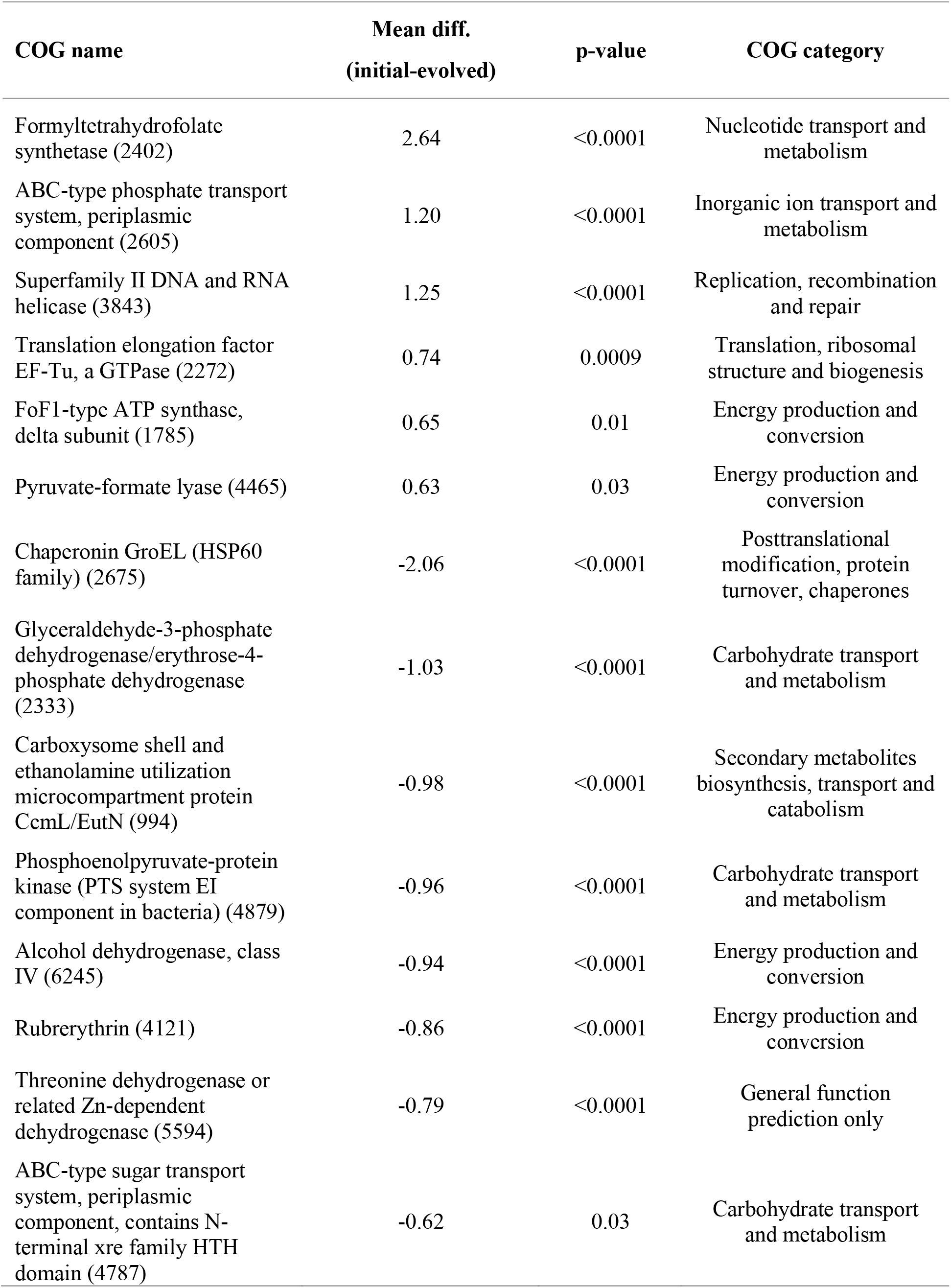
Fourteen COGs significantly different between initial and evolved samples.

**FIGURE 5.**
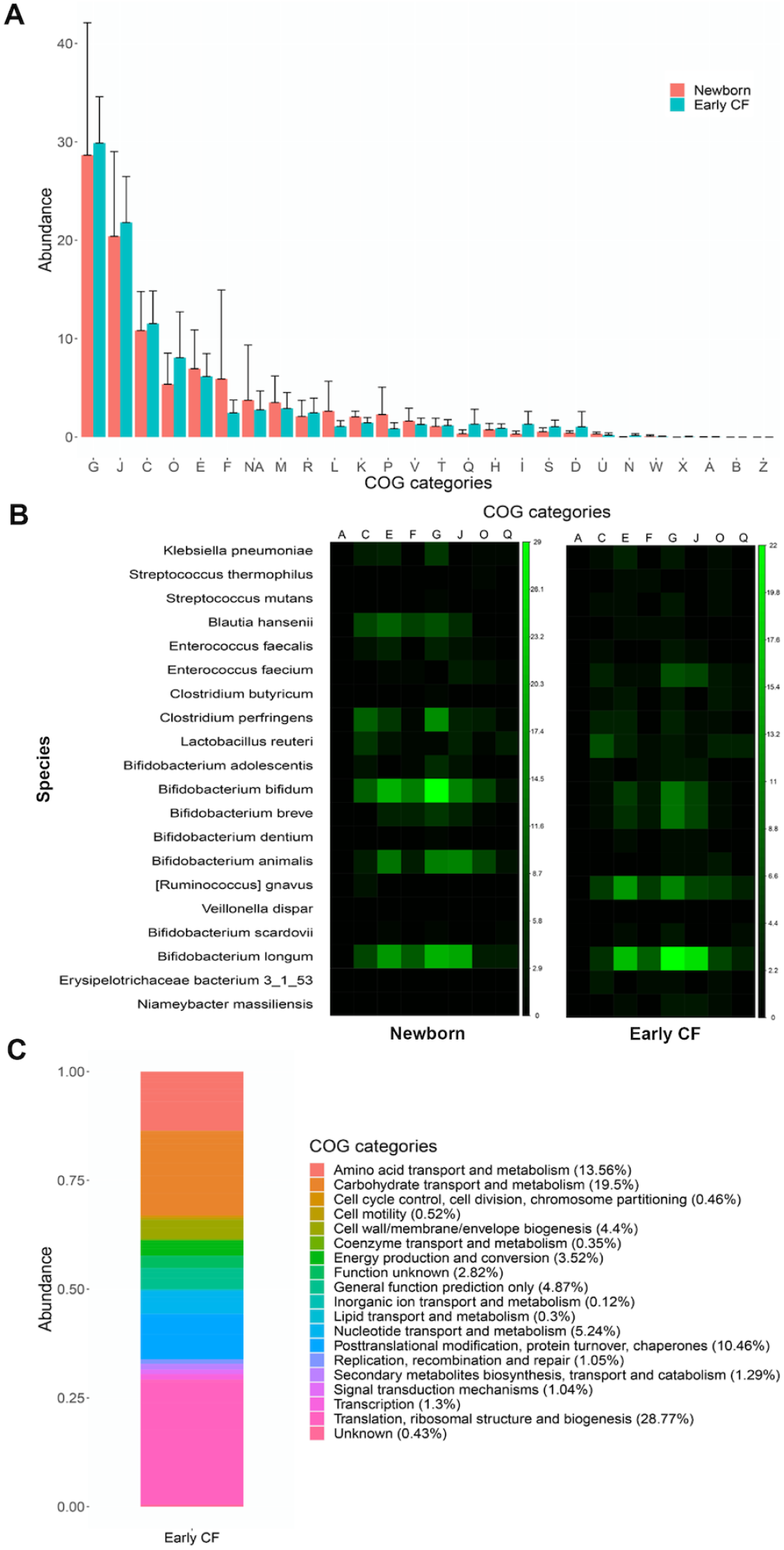
**A)** Bar plot representing the different COG categories in the newborn and early CF groups. The abundance is represented as the mean percentage of the sum of all the peptides correlated with this COG category/total sum of the peptide’s intensity of each group (newborn or early CF). **B)** Function and taxon correlation analysis at species level of the proteins detected in newborn and early CF groups. The color scale represents the intensity of the proteins associated to the specific species in the different COG categories. **C)** Histogram of the COG category distribution of the 69 proteins detected from *R. gnavus* in early CF group. The abundance of the proteins is the sum of peptides intensity belonging to that protein. The percentage is calculated as the sum of the intensity of the proteins associated to that COG category/sum of the total intensities of proteins from *R. gnavus*. COG categories nomenclature: A, RNA processing and modification; B, Chromatin structure and dynamics; C, Energy production and conversion; D, Cell cycle control, cell division, chromosome partitioning; E, Amino acid transport and metabolism; F, Nucleotide transport and metabolism; G, Carbohydrate transport and metabolism; H, Coenzyme transport and metabolism; I, Lipid transport and metabolism; J, Translation, ribosomal structure and biogenesis; **K**, Transcription; **L**, Replication, recombination and repair; **M**, Cell wall/membrane/envelope biogenesis; **N**, Cell motility; O, Posttranslational modification, protein turnover, chaperones; P, Inorganic ion transport and metabolism; Q, Secondary metabolites biosynthesis, transport and catabolism; R, General function prediction only; S, Function unknown; T, Signal transduction mechanisms; U, Intracellular trafficking, secretion, and vesicular transport; V, Defense mechanisms; W, Extracellular structures; X, Mobilome: prophages, transposons; Z, Cytoskeleton; NA, not assigned to any COG category

The correlation of each bacterial taxa and their function was also performed with iMetaLab, using the iMetaShiny Apps. *Bifidobacterium spp*. was the most abundant taxa and was also responsible for most of the bacterial functions in the initial samples (Figure 5B). Bifidobacterial proteins were still abundant in the evolved samples, in which most of the functional pathways were promoted by *R. gnavus* (Figure 5B). A significant enrichment of *R. gnavus* proteins was observed from baseline to evolved times (2 vs. 69 proteins), most of them related to translational processes, such as amino acid and carbohydrate metabolism (Figure 5C). In contrast, the functions associated with *Clostridium perfringens* and *Blautia hansenii* showed a statistically significant decrease in the evolved samples.

Metaproteomics also allowed the identification of 293 human proteins: 85 exclusively present in baseline samples, 43 exclusively in evolved samples, and 165 in both samples (Additional File 6). Six of the 165 shared proteins had significantly differential expression levels: lactotransferrin (*p*<0.0001) and myeloperoxidase (*p*<0.0001) were highly expressed in the newborn samples, whereas actin, cytoplasmic 2 (*p*<0.0001), chymotrypsin-C (*p*<0.0001), carboxypeptidase A1 (*p=*0.0409), and myosin-1 (*p*<0.0001) were in higher abundance in the most evolved CF samples. A STRING analysis (STRING version 11.5) was performed with the 3 groups of human proteins (newborn, early CF, and shared proteins) including for the first 2 groups those proteins exclusively identified and with a significantly higher abundance in each of the samples (Figure 6). The analysis of the proteins shared by the two groups showed a significant protein-protein interaction (1.87 × 10^−16^). Shared proteins were mainly related to neutrophil-mediated immunity (42 proteins, *p*=1.87 × 10^−25^), such as calprotectin (S100-A9 and S100-A8), neutrophil elastase (ELANE), cathepsin G, and myeloperoxidase, or related to the intestinal barrier, such as mucins (MUC2, MUC4, MUC13) or proteins present in cell junctions (40 proteins, *p*=1.24 × 10^−5^) (Figure 6A). The initial samples contained significantly more proteins associated with chylomicron assembly and the immune system than the final samples (Figure 6B), in which a cluster of muscle proteins stands out (Figure 6C). The names of the proteins belonging to each function are listed in Additional File 7.

**FIGURE 6.**
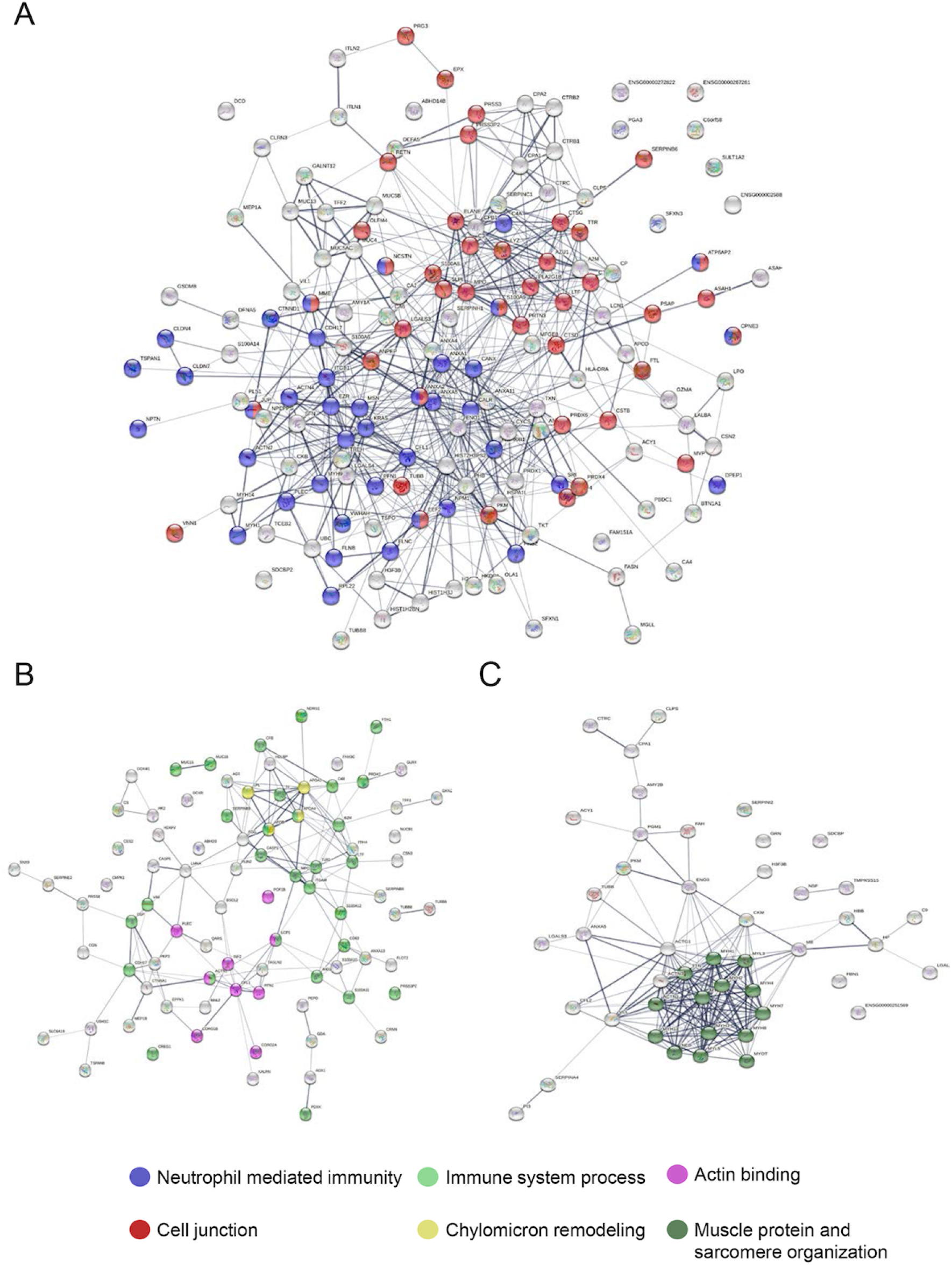
STRING map protein-protein interaction network of the human proteins detected in this work in A) shared proteins between newborn and early CF groups, B) newborn group (including proteins exclusively identified in newborn samples and those more abundant in this group) and C) early CF group (including proteins exclusively identified in early CF samples and those more abundant in this group). Line thickness indicates the strength of data support. The colors represent different processes.

## DISCUSSION

A major clinical feature of CF are digestive comorbidities, such as steatorrhea, maldigestion/malabsorption, and pancreatic insufficiency, also affect the quality of life and nutritional status of these individuals. The CF environment strongly conditions the gut microbiota composition (31), also contributing to their malfunction, even at the age of 6 weeks (9). Even though amplicon sequencing is the most widely used technique for determining the composition of gut microbiota, there is a growing demand for techniques that determine the global functionality of the ecosystem, such as metaproteomics. This study aimed to monitor the establishment of the CF gut microbiota during the first year of life, using amplicon sequencing and metaproteomics and incorporating a robust statistical analysis to compare their level of agreement in taxonomic assignment and abundance detection.

The taxonomical incongruities detected were attributed to paraphyletic groups, which are continuously reclassified according to the latest advances in phylogeny. In this sense, SILVA 138 is a more accurate database with which to establish phylogenetic relationships, because protein databases are less frequently updated. Of more concern was the exclusive detection of certain taxa by one of the two methodologies, particularly at the species level.

Curiously, metaproteomics detected proteins attributed to *Streptococcus thermophilus*, whereas when using amplicon sequencing, the proteins appeared to correspond to *Streptococcus salivarius*. Poor resolution of streptococcal species using the V3-V4 regions of the 16S rRNA gene has been previously reported (32). Another amplicon sequencing limitation is the detection of what is known as sporobiota (33), most of the endosporulating species belonging to *Firmicutes*. Spores are underrepresented because of their ability to resist DNA extraction techniques, the high similarity between 16S rRNA and housekeeping genes of other sporulating bacteria, and their larger genomes that, when sequenced, produce fewer reads per gene per taxon. Other bacterial phyla, such as *Actinobacteria* (particularly *Bifidobacterium*), have genomes with a high GC content, and this likewise contributes to their underestimation by sequencing techniques.

Metaproteomics has also failed in detecting or properly estimating certain bacterial groups typically found in human gut microbiota due to their resistance to cell lysis, especially Gram-positive bacteria (34). Moreover, certain microbial components and detergents used for cell lysis can affect post-enzymatic digestion (35). Our workflow includes optimized steps, such as serial centrifugations for microbial pre-enrichment, sonication with bead beating, and lysis with SDS-based buffers. Whereas our method improves the number of peptide and protein identifications and the relative abundance of *Actinobacteria*, other protocols have shown a higher identification of proteins from other taxonomic groups, such as *Proteobacteria*, but a lower number of peptide and protein identifications (36). Metaproteomic analysis presents several bioinformatics challenges, the most common being adequate peptide identification, taxonomic assignment, and the quality of functional annotation (35). It is estimated that only 20% of identified peptides reach the species level.

The statistical analysis of our results demonstrated an equivalent bacterial taxonomic assignation by both techniques, while for abundance detection some discrepancies have found by the previously used Bland-Altman test (37). The underrepresentation of *Bifidobacterium* by amplicon sequencing has been repeatedly notified, but by the genomic technique we also detected an overestimation of some *Firmicutes* and *Proteobacteria* abundance (38).

The alpha diversity indexes of our CF infant cohort did not reach those observed in other CF cohorts nor in the control cohort by Antosca *et al*. (9), although other authors have reported lower values than for healthy controls (39). A discrete increase was observed in the evolved CF samples, probably as a direct consequence of antibiotic intake.

Fecal proteomics in adults with CF have pointed to the dominance of *R. gnavus, Enterobacteriaceae*, and *Clostridia* species, combined with decreased butyrate producers, such as *Faecalibacterium prausnitzii* (17). We detected similar compositional changes in a very reduced cohort of 8 infants followed for almost 18 months right after being diagnosed with CF. *R. gnavus* has been associated with several human gastrointestinal diseases and is a mucolytic bacterium capable of degrading the human colonic mucin, MUC2, which is a major form of secretory glycosylated mucin coating the epithelia of the intestine and therefore has a close relationship with the mucus layer (40, 41). It has been suggested that the bacterium can penetrate the mucus layer and lead to an activation of the immune system and inflammatory responses (42, 43). Nevertheless, the involvement of this species in allergic or autoimmune diseases has been also questioned (40, 44).

Drastic changes in the bacterial functional profile when comparing the initial and evolved samples were not observed; the proteins more abundant in the initial samples were a formyltetrahydrofolate synthetase and a pyruvate-formate lyase previously related to biofilm upregulation (45). In the final samples, the most relevant enrichment was the chaperonin GroEL, a stress response protein whose main role is protein folding, which has recently been observed to have roles in virulence and pathogenesis, such as bacterial adherence and immune invasion (46).

One of the most interesting advantages of metaproteomics is the detection of human proteins. In our samples, several proteins associated with the maintenance of the intestinal epithelium and the immunity response were detected. Lactotransferrin, the iron sequestration protein abundant in human breast milk, and myeloperoxidase, inflammation regulated by granulocytes of neutrophils, were significantly enriched in the initial samples (47, 48). All our results point to an altered establishment of the gut microbiota in infants with CF, with a dominance of *R. gnavus* within a human inflammatory state.

Our study provides an extended comparative analysis with robust statistical support that could optimize the use of both approaches for gut microbiota research. Metaproteomics provides information on composition and functionality, as well as data on host-microbiome interactions. Its strength is the identification and quantification of *Actinobacteria* and certain classes of *Firmicutes*. Taking all the results into account, both techniques detected an aberrant microbiota in infants with CF during their first year of life, dominated by the enrichment of *R. gnavus* within a human inflammatory environment.

## MATERIALS AND METHODS

### Participants, sample collection, and medical records

Subjects included in this study had CF diagnosed by neonatal screening, sweat chloride test, and mutation sequencing, and recruited during their first month of life, when the first fecal sample was collected, between January 2018 and January 2019. The infants were attended at 3 Spanish CF reference hospitals, geographically distant. Ethical approval for the study was granted by the Ramón y Cajal Ethics Committee in 2017, and all of the infants’ parents signed the informed consent. Subjects were excluded if they were born after that date, had less than 3 collected samples, or if important clinical data was missing. Eight infants with CF were finally followed up, and the relevant data, including feeding habits, delivery mode, and CF mutations of them are shown in Table 1. Data related to antibiotic and other treatments’ usage is summarized in Additional File 1. Ethical approval for the study was granted by the Ramón y Cajal Ethics Committee in 2017, and all of the infants’ parents signed the informed consent. Fecal samples were sequentially collected from the diaper of each infant during the first year of life over routinary medical chek-ups. Samples were stored at −80°C in 2 aliquots for subsequent amplicon sequencing and metaproteomic analysis.

### DNA extraction and 16S rRNA amplicon sequencing

Our study followed guidelines for the Strengthening The Organization and Reporting of Microbiome Studies (STORMS) reporting (Additional File 2) (22). Total DNA from the fecal samples was obtained with the QiaAMP kit (Qiagen, Germany), and further Mi-Seq 2×300 bp paired-end (Illumina) 16S rRNA sequencing of the V3 and V4 regions (23) was performed at the Central Unit for Translational Genomics Support (Ramon y Cajal Health Research Institute). The sequencing data analysis was performed using the Qiime pipeline (24), which includes DADA2 for sequence quality filtering, and a Silva 138 database (released: Dec 2019) was used for taxonomical assignment, discharging those samples with fewer than 1000 reads (n=2). Nucleotide sequences were deposited in the National Center for Biotechnology Information’s Sequence Read Archive repository, BioProject ID 719717 (https://www.ncbi.nlm.nih.gov/bioproject/PRJNA719717).

### Metaproteomics

Samples were prepared using differential centrifugation, and the microbial pellet was processed by sonication according to a previously published protocol. In brief, 0.1-0.3 g of feces were suspended in 10 mL phosphate-buffered saline (PBS) and mixed in a tube rotator for 45 min at 4 ºC. The samples were centrifuged at 500 g for 5 min, and the supernatant was collected in a 50-mL tube. Ten mL of PBS was added again, repeating the process twice, and the 3 supernatants (∼30 mL) were centrifuged at 11,000 g. Microbial pellets were suspended in 500 µL of lysis buffer (4% sodium dodecyl sulfate [SDS], 50 mM Tris-HCl pH 8.0) and heated for 10 min at 95 ºC. Four sonication cycles (30 s with a 1-min interval on ice) were performed, with an amplitude of 40%. Silica beads were then added (0.3 g) to each sample, and 5 rounds of bead beating were performed (30 s with a 5-min interval on ice) at a speed of 6.5 ms^-1^. A 14,000-g centrifugation was performed to remove the beads and cell debris. To remove the SDS, proteins were precipitated using methanol/chloroform and suspended in 8 M urea for in-solution trypsin digestion. Lastly, the peptides were quantified in a Qubit fluorimeter (Thermo Scientific), and 1 µg of peptides were loaded for reverse-phase-nano-liquid chromatography electrospray ionization tandem mass spectrometric analysis on an EASY-nLC 1000 System (Proxeon) coupled to a Q-Exactive HF mass spectrometer (Thermo Scientific Inc.). Peptides were loaded on-line onto an Acclaim PepMap 100 Trapping column (75 μm i.d. × 20 mm, 3 μm C18 resin with 100 Å pore; Thermo Scientific) using buffer A (0.1% formic acid) and then separated on a C18 resin analytical column (75 μm i.d. × 500 mm, 2 µm and 100 Å pore size; Thermo Scientific Easy Spray Column). A 240-min gradient from 2% to 40% buffer B (0.1% formic acid in 100% acetonitrile) in buffer A was performed to separate the peptides.

Data were obtained by data-dependent acquisition in positive mode. From each MS scan (between 350 and 2000 Da), the 15 most intense precursors (charge between 2+ and 5+) were selected for their high collision energy dissociation fragmentation, with a dynamic exclusion of 10 s and a normalized collision energy of 20, and the corresponding MS/MS spectra were acquired. Peptides were eluted using a 240 min gradient. The MS proteomics raw data have been submitted to the ProteomeXchange Consortium (http://www.proteomexchange.org) via the Proteomics Identifications Database partner repository with the database identifier PXD029284 (25). For data processing, we employed MetaLab software, which provides a human gut microbial database (12, 26). A human database downloaded from Uniprot DB (http://www.uniprot.org) (27), restricted to human taxonomy (downloaded on 02/18/2020 with 74,451 sequences), was also used to identify human proteins. For peptide and protein identification, the false discovery rate was set to 0.01. Only taxa identified with at least 2 peptides were considered and were manually filtered to eliminate human peptides. The sum of the intensities of all the distinctive peptides assigned to a taxon was used as the relative abundance of that taxon. A taxon-function analysis was also performed, using taxonomic information of the enrichment analysis from the iMetaLab platform (http://shiny.imetalab.ca/) (28). GraphPad Prism (version 9.1.1.225) was used for the functional statistical analysis of the microbial and human proteins. A Search Tool for the Retrieval of Interacting Genes/Proteins (STRING) analysis was performed with the STRING program (version 11.5). Graphs were constructed with Rstudio (version 1.4.1717).

### Taxonomic assignment and microbial quantification agreements

The data were trimmed by selecting only the taxa identified by both methods (shared taxa), solving annotation incongruences. First, we evaluated the dichotomous dependent variable, “taxa presence,” by a McNemar test with continuity correction at the phylum, class, and genus levels, considering in each sample any operational taxonomic unit (OTU) or at least 2 peptides per taxa.

Second, the relative abundance of each phylum, class, or genus was normalized per sample, expressed as a percentage, and the level of agreement between the two techniques was assessed by Bland–Altman plots, representing the mean of relative abundances (%) against the difference of relative abundances (%) per sample (amplicon sequencing minus metaproteomics abundance), including an upper and lower limit of agreement calculated as the mean difference ±1.96-fold the standard deviation. Only the taxa detected in more than 50% of the samples by at least one of the approaches were included for this abundance analysis. A nonparametric Wilcoxon rank-sum paired test was performed to compare the populations’ mean ranks and contribute to the descriptive plots.

### Gut microbiota establishment in CF

Differences between each infant’s first and last fecal samples in terms of bacterial composition and distribution were evaluated, employing the phyloseq R package (29) using normalized results per sample: only OTUs or amplicon sequence variants (ASVs) with more than 2 counts and that were present in more than 10% of samples were preserved. Alpha diversity was analyzed by the Shannon–Weaver diversity index, and differences were evaluated by the Mann– Whitney/Kruskal–Wallis statistical method. Differential abundance in certain taxa above a fold-change threshold (variance in abundance of 10^5^ for sequencing data and 10^8^ for proteomics data) was evaluated using the edgeR package in RStudio (30).

## Data availability

The sequenced data from this article is deposited in the NCBI’s Sequence Read Archive (SRA) repository, BioProject ID 719717 (https://www.ncbi.nlm.nih.gov/bioproject/PRJNA719717). The proteomics data set from this paper has been deposited in the ProteomeXchange Consortium via the PRIDE partner repository with the data set identifier PXD029284.

## ACKNOWLEDGMENTS

We thank the families of the children for their participation and all the health personnel of the CF Unit, and Marta Cobo for technical assistance.

